# Vitamin B2 enables peroxisome proliferator-activated receptor α regulation of fasting glucose availability

**DOI:** 10.1101/2022.10.16.512439

**Authors:** Peter M. Masschelin, Pradip K. Saha, Scott A. Ochsner, Aaron R. Cox, Kang Ho Kim, Jessica B. Felix, Robert Sharp, Xin Li, Lin Tan, Jun Hyoung Park, Liping Wang, Vasanta Putluri, Philip L. Lorenzi, Alli M. Nuotio-Antar, Zheng Sun, Benny Kaipparettu, Nagireddy Putluri, David D. Moore, Scott A. Summers, Neil J. McKenna, Sean M. Hartig

## Abstract

Flavin adenine dinucleotide (FAD) interacts with flavoproteins to mediate oxidation-reduction reactions required for cellular energy demands. Not surprisingly, mutations that alter FAD binding to flavoproteins cause rare inborn errors of metabolism (IEMs) that disrupt liver function and render fasting intolerance, hepatic steatosis, and lipodystrophy. In our study, depleting FAD pools in mice with a vitamin B2 deficient diet (B2D) caused phenotypes associated with organic acidemias and other IEMs, including reduced body weight, hypoglycemia, and fatty liver disease. Integrated discovery approaches revealed B2D tempered fasting activation of target genes for the nuclear receptor PPARα, including those required for gluconeogenesis. Treatment with the PPARα agonist fenofibrate activated the integrated stress response and refilled amino acid substrates to rescue fasting glucose availability and overcome B2D phenotypes. Overall, these findings reveal PPARα governs metabolic responses to FAD availability and nominate its pharmacologic activation as strategies for organic acidemias.

## Introduction

Flavin mononucleotide (FMN) and flavin adenine dinucleotide (FAD) serve as essential cofactors for diverse proteins that mediate oxidation-reduction (redox) reactions, transcriptional regulation, and metabolism (**Powers, 2003**). In particular, FAD supports the activity of flavoproteins that enable the electron transport chain (ETC), the tricarboxylic acid (TCA) cycle, and fatty acid oxidation (FAO). Along these lines, mutations occurring in more than 50% of human flavoproteins cause inborn errors of metabolism (IEMs) with heterogeneous clinical presentations frequently characterized by organic acidemia, fasting intolerance, and fatty liver disease (**Balasubramaniam et al., 2019**). Compared to the more well understood roles of nicotinamide adenine dinucleotide (NAD), the physiological relevance of FAD has remained largely ignored. This gap in knowledge slows the pursuit of therapeutic strategies to treat IEM and leaves fundamental energy balance roles for FAD unresolved.

The liver coordinates whole-body metabolism during fasting by releasing glucose and other fuels to spare the brain and survive low-nutrient conditions. Nuclear receptors sense and receive the signals of nutrient abundance in the liver to perform precise regulation of genes that ultimately maintain the energetic needs of the FAO and amino acid catabolism pathways serving gluconeogenesis (**Scholtes and Giguère, 2022**). Among nuclear receptors, peroxisome proliferator-activated receptor a (PPARα) modulates fasting responses and pivots hepatocytes towards conservation and recycled substrates to sustain energy production. Synthetic PPARα agonists promote FAO by directing the activity of pathways for balanced lipid metabolism in the liver, which consequently supported a series of FDA-approved fibrate drugs for the treatment of hypertriglyceridemia (**Jackevicius et al., 2011**). The lipid-lowering properties of bezafibrate and fenofibrate motivated pre-clinical studies (**Steele et al., 2020; Waskowicz et al., 2019; Yavarow et al., 2020**) that form ongoing efforts to overcome the limited armament of therapies for IEMs.

Nutrition impacts FAD availability for the organism and dietary riboflavin (vitamin B2) supplies the backbone for all FAD and FMN synthesis. Thus, riboflavin deficiency gives rise to abnormal development and energy balance disorders. Models that expand how FAD requirements form the regulatory environment for metabolic homeostasis provide opportunities for pre-clinical experiments and studies of fundamental nutrient-sensing mechanisms. Here, we define outcomes of vitamin B2 depletion resembling IEMs of flavoprotein deficiency and transcriptional pathways that preserve glucose availability in fasted mice.

## Materials and methods

### Mice and housing conditions

All animal procedures were approved by the Institutional Animal Care and Use Committee of Baylor College of Medicine (AN-6411). All mice were housed in a barrier-specific pathogen-free animal facility with 12 h dark-light cycle and free access to water and food. C57BL/6J wild-type mice (RRID:IMSR_JAX:000664) were obtained from the BCM Center for Comparative Medicine and global *Ppara^-/-^* mice (RRID:IMSR_JAX:008154) were generated previously (**Lee et al., 1995**). In all experiments, male mice were randomly placed on control or riboflavin deficient diet starting at 4-weeks of age. Riboflavin-deficient and matched control diets were provided by Research Diets: 90% Control, D10012G; 90% Riboflavin-Deficient, D12030102; 99%, Control A18041301; or 99% Riboflavin-Deficient, A19080901. All diets were isocaloric, and amino acids were kept constant **(Supplemental Table S1 & S2)**. All experiments adhered to ARRIVE guidelines.

### Fenofibrate gavage

A 0.5% methylcellulose solution was prepared by heating 150mL water with 2.5g 400cP methylcellulose (Sigma, #M0262) added with stirring. Chilled water was added (350ml) and stirred overnight at 4°C. Fresh fenofibrate-gavage solution was made daily with 112.5mg fenofibrate (Sigma, #F6020) in 1.5mL 0.5% methylcellulose solution. Mice were gavaged daily at 300mg/kg.

### Pyruvate tolerance test (PTT)

To determine pyruvate tolerance, mice were fasted for 16h, and Na-pyruvate was administered (1g/kg body weight) by intraperitoneal injection. Blood glucose levels were monitored at 0, 15, 30, 60, and 120 minutes by a Freestyle Glucose Monitoring System (Abbott Laboratories).

### Glucose production rate

In *vivo* glucose production was performed, beginning with the insertion of a microcatheter into the jugular vein under anesthesia, followed by 4-5 days rest for complete recovery. Overnight-fasted (16h) conscious mice received a priming bolus of HPLC-purified [3-^3^H] glucose (10μCi) and then a constant infusion (0.1μCi/min) of labeled glucose for ~90 minutes. Blood samples were collected from the tail vein at 0, 50, 60, 75, and 90 min to calculate the basal glucose production rate from tracer dilution. Steady states were reached within one hour of infusion.

### Indirect calorimetry

Wild-type mice were maintained on experimental diets and housed at room temperature in Comprehensive Lab Animal Monitoring System Home Cages (CLAMS-HC, Columbus Instruments). Oxygen consumption, CO_2_ emission, energy expenditure, food and water intake, and activity were measured for five days (BCM Mouse Metabolic and Phenotyping Core). Mouse body composition was examined by magnetic resonance imaging (Echo Medical Systems) before indirect calorimetry.

### Histology

Sections of liver tissue were frozen in Tissue-Tek OCT compound (4583; Sakura Finetek USA), and neutral lipids stained with Oil Red O. Formalin-fixed paraffin-embedded liver tissue sections were stained with hematoxylin and eosin (H/E). Images were captured on a Nikon Ci-L Brightfield microscope.

### Serum and lipid assays

Fasted serum was used to measure serum lactate (K607; Biovision), ALT (TR71121; Thermo Scientific), AST (TR70121; Thermo Scientific), beta-hydroxybutyrate (Biovision K632), and serum-free fatty acids (#sfa-1; Zen-Bio).

### Liver FAD and glycogen measurements

10-15 mg of liver tissue was deproteinated (K808; Biovision), followed by measurement of FAD through a colorimetric assay (K357; Biovision). FAD concentration was standardized to input tissue weight. 10-15 mg of liver tissue was used for measuring fasting hepatic glycogen (K646; Biovision).

### Hepatic TG and cholesterol

Both serum and tissue samples were analyzed for triglycerides (Triglyceride reagent TR22421; Thermo Scientific) and cholesterol (Total Cholesterol Reagent TR13421; Thermo Scientific). Hepatic TGs and cholesterol were assayed as described previously (**Kim et al., 2019**). Briefly, liver homogenates were mixed with a 1:2 chloroform:methanol solution followed by isolation of the lipid-rich chloroform layer (modified Folch method).

### Immunoblotting

Cell and tissue lysates were prepared in Protein Extraction Reagent (ThermoFisher) supplemented with Halt Protease and Phosphatase Inhibitor Cocktail (ThermoFisher). Western blotting was performed with whole-cell lysates run on 4-12% Bis-Tris NuPage gels (Life Technologies) and transferred onto Immobilon-P Transfer Membranes (Millipore), followed by antibody incubation. Immunoreactive bands were visualized by chemiluminescence. Antibodies used in this study are listed in **Supporting Table S5**.

### RNA extraction and RNA-seq analysis

Total liver RNA was extracted using the Qiagen RNeasy Plus Mini kit (74034; Qiagen). Sample quality was confirmed on an Agilent 2100 Bioanalyzer (Agilent). mRNA library preparation and RNA sequencing were performed by Novogene. mRNA libraries were prepared with NEBNext Ultra RNA Library Prep Kit for Illumina (NEB) and size selection for libraries was performed using AMPure XP system (Beckman Coulter), followed by library purity analysis. Libraries were sequenced on NovaSeq PE150 and reads mapped to the UCSC mouse reference genome mm10 using STAR. FeatureCounts was used to calculate the expression level as reads per kilobase per million (RPKM). DESeq2 calculated differentially expressed genes with p values adjusted using Benjamini and Hochberg’s method for controlling the False Discovery Rate (FDR). Genes with significant differential expression were determined by p < 0.05. Gene set enrichment analysis was performed with the Molecular Signatures Database, and −log_10_(p-value) calculated for Hallmark gene sets.

### Consensome and high confidence transcriptional (HCT) target intersections

Transcription factor footprint analysis and consensome enrichments were performed as previously described (**Ochsner et al., 2019**). For transcription factors, node and node family consensomes are gene lists ranked according to the strength of their regulatory relationship with upstream signaling pathway nodes derived from independent publicly archived transcriptomic or ChIP-Seq datasets. Genes in the 95th percentile of a given node consensome were designated high confidence transcriptional targets (HCTs) for that node and used as the input for the HCT intersection analysis using the Bioconductor GeneOverlap analysis package implemented in R. For both consensome and HCT intersection analysis, p values were adjusted for multiple testing using the method of Benjamini and Hochberg to control the FDR as implemented with the p. adjust function in R, to generate q values. Evidence for a transcriptional regulatory relationship between a node and a gene set was inferred from a larger intersection between the gene set and HCTs for a given node or node family than would be expected by chance after FDR correction (q < 0.05). The HCT intersection analysis code has been deposited in the SPP GitHub account at https://github.com/signaling-pathways-project/ominer/.

### Metabolomics

Targeted measurement of hepatic carnitines, fatty acids, lipids species, CoA’s, glycolysis, and TCA metabolites was carried out by the BCM Dan L Duncan Cancer Center CPRIT Cancer Proteomics and Metabolomics Core. Parallel analysis of lipids and ceramides was performed at the University of Utah. All samples were processed and analyzed as described previously (**Chaurasia et al., 2019; Kettner et al., 2016**).

#### Reagents

High-performance liquid chromatography grade and mass spectrometry grade reagents were used: acetonitrile, methanol, and water (Burdick & Jackson); formic acid, ammonium acetate, and internal standards (Sigma-Aldrich); MS grade lipid standards (Avanti Polar Lipids).

#### Internal Standards and Quality Control

To assess overall process reproducibility, mouse pooled liver or serum samples were run along with the experimental samples. A number of internal standards, including injection standards, process standards, and alignment standards, were used to assure QA/QC targets to control for experimental variability. Aliquots (200μL) of 10mM solutions of isotopically labeled standards were mixed and diluted up to 8000μL (final concentration 0.25mM) and aliquoted into a final volume of 20μL. The aliquots were dried and stored at −80°C until further analysis. To monitor instrument performance, 20μL of a matrix-free mixture of the internal standards were reconstituted in 100μL of methanol:water (50:50) and analyzed by MRM. The metabolite extraction from the samples was monitored using pooled mouse serum or liver samples and spiked internal standards. The matrix-free internal standards and serum and liver samples were analyzed twice daily. The median coefficient of variation (CV) value for the internal standard compounds was 5%. To address overall process variability, metabolomic studies were augmented to include a set of nine experimental sample technical replicates, which were spaced evenly among the injections for each day.

#### Separation of CoAs and carnitines

Targeted profiling for CoAs and carnitines in electro spray ionization positive mode by the RP chromatographic method employed a gradient containing water (solvent A) and acetonitrile (ACN, solvent B, with both solvents containing 0.1% formic acid). Separation of metabolites was performed on a Zorbax Eclipse XDBC18 column (50 × 4.6mm i.d.; 1.8μm, Agilent Technologies) maintained at 37°C. The gradient conditions were 0-6 minutes in 2%B; 6.5 minutes in 30% B, 7 minutes in 90% B, 12 minutes in 95% B, followed by re-equilibration to the initial conditions.

#### Separation of glycolysis and TCA metabolites

The glycolysis and TCA metabolites were separated by the normal phase chromatography using solvents containing water (solvent A), solvent A modified by the addition of 5mM Ammonium acetate (pH 9.9), and 100% acetonitrile (solvent B). The binary pump flow rate was 0.2mL/min with a gradient spanning 80% B to 2% B over 20 minutes, 2% B to 80% B for 5 minutes, and 80% B for 13 minutes. The flow rate was gradually increased during the separation from 0.2mL/min (0-20 minutes) to 0.3mL/min (20-25 minutes), and then 0.35mL/min (25-30 minutes), 0.4mL/min (30-37.99 minutes) and finally to 0.2mL/min (5 minutes). Metabolites were separated on a Luna Amino (NH2) column (4μm, 100A 2.1×150mm, Phenominex) maintained in a temperature-controlled chamber (37°C). All the columns used in this study were washed and reconditioned after every 50 injections.

#### Separation of fatty acids

Targeted profiling for fatty acids employed the RP chromatographic method by a gradient containing water (solvent A) with 10 mM ammonium acetate (pH 8) and 100% methanol (solvent B) on a Luna Phyenyl Hexyl column (3μm, 2X150mm; Phenominex, CA) maintained at 40°C. The binary pump flow rate was 0.2 mL/min with a gradient spanning 40% B to 50% B over 8 minutes, 50% B to 67% B over 5 minutes, hold 67% B for 9 minutes, 67% B to 100% B over 1 minute, hold 100% B for 6 minutes, 100% B to 40% B over 1 minute and hold 40% B for 7 minutes.

#### Liquid chromatography/mass spectrometry (LC/MS) analyses

The chromatographic separation of non-lipid metabolites was performed using either reverse phase separation or normal phase online with the unbiased profiling platform based on a 1290 SL Rapid resolution LC and a 6490 triple Quadrupole mass spectrometer (Agilent Technologies, Santa Clara, CA). Lipidomics required a Shimadzu CTO-20A Nexera X2 UHPLC coupled with TripleTOF 5600 equipped with a Turbo VTM ion source (AB Sciex, Concord, Canada). Using a dual electrospray ionization source, the samples were independently examined in both positive and negative ionization modes. The data acquisition during the analysis was controlled using the Mass Hunter workstation data acquisition software.

#### Lipidomics

Mouse liver lipids were extracted using a modified Bligh-Dyer method. Briefly, 50 mg of crushed tissue sample from mouse whole liver was used. A 2:2:2 volume ratio of water/methanol/dichloromethane was used for lipid extract at room temperature after spiking internal standards 17:0 LPC, 17:0PC, 17:0 PE, 17:0 PG, 17:0 ceramide, 17:0 SM, 17:0PS, 17:0PA, 17:0 TAG, 17:0MAG, DAG 16:0/18:1, CE 17:0. The organic layer was collected, followed by a complete drying procedure under nitrogen. Before MS analysis, the dried extract was resuspended in 100μL of Buffer B (10:5:85 Acetonitrile/water/Isopropyl alcohol) containing 10mM NH4OAc and subjected to reverse-phase chromatography and LC/MS. Internal standards prepared in chloroform/methanol/water (100pmol/μL) were LPC 17:0/0:0, PG 17:0/17:0, PE 17:0/17:0, PC 17:0/17:0, TAG 17:0/17:0/17:0, SM 18:1/17:0, MAG 17:0, DAG 16:0/18:1, CE 17:0, ceramide 18:1/17:0, PA 17:0, PI 17:0/20:4, and PS 17:0/17:0.

For lipid separation, 5mL of the lipid extract was injected into a 1.8 mm particle 50 × 2.1mm Acquity HSS UPLC T3 column (Waters). The column heater temperature was set at 55°C. For chromatographic elution, a linear gradient was used over a 20 min total run time, with 60% Solvent A (acetonitrile/water (40:60, v/v) with 10mM ammonium acetate) and 40% Solvent B (acetonitrile/water/isopropanol (10:5:85 v/v) with 10mM ammonium acetate) gradient in the first 10 min. The gradient was ramped linearly to 100% Solvent B for 7 min. Then the system was switched back to 60% Solvent B and 40% Solvent A for 3 min. A 0.4mL/min flow rate was used at an injection volume of 5μL. The column was equilibrated for 3 min and run at a flow rate of 0.4mL/min for a total run time of 20 min. TOF MS survey scans (150ms) and 15 MS/MS scans with a total duty cycle time of 2.4s were performed. The mass range in both modes was 50-1200m/z. The acquisition of MS/MS spectra by data-dependent acquisition (DDA) function of the Analyst TF software (AB Sciex).

The raw data in .mgf format was converted using proteoWizard software. The NIST MS PepSearch Program was used to search the converted files against LipidBlast libraries. The m/z width was determined via the mass accuracy of internal standards at 0.001 for positive mode and 0.005 for a negative mode at an overall mass error of less than 2 ppm. The minimum match factor at 400 was set for the downstream data processing. The MS/MS identification results from all the files were combined using an in-house software tool to create a library for quantification. The raw data files were searched against this in-house library of known lipids with mass and retention time using Multiquant 1.1.0.26 (ABsciex). The lipid species identified in the positive or negative ion modes were analyzed separately using relative abundance of peak spectra for the downstream analyses. The identified lipids were quantified by normalizing against their respective internal standard.

#### Ceramides and lipids

Lipid extracts are separated on an Acquity UPLC CSH C18 1.7μm 2.1 × 50mm column maintained at 60°C connected to an Agilent HiP 1290 Sampler, Agilent 1290 Infinity pump, equipped with an Agilent 1290 Flex Cube and Agilent 6490 triple quadrupole (QQQ) mass spectrometer. In positive ion mode, sphingolipids are detected using dynamic multiple reaction monitoring (dMRM). Source gas temperature is set to 210°C, with a N2 flow of 11L/min and a nebulizer pressure of 30psi. Sheath gas temperature is 400°C, sheath gas (N2) flow of 12L/min, capillary voltage is 4000V, nozzle voltage 500V, high-pressure RF 190V, and low-pressure RF is 120V. Injection volume is 2μL, and the samples are analyzed in a randomized order with the pooled QC sample injection eight times throughout the sample queue. Mobile phase A consists of ACN: H2O (60:40 v/v) in 10mM ammonium formate and 0.1% formic acid, and mobile phase B consists of IPA: ACN:H2O (90:9:1 v/v) in 10mM ammonium formate and 0.1% formic acid. The 5 chromatography gradient starts at 15% mobile phase B, increases to 30% B over 1 min, increases to 60% B from 1-2 min, increases to 80% B from 2-10 min, and increases to 99% B from 10-10.2 min where it’s held until 14 min. Post-time is 5 min, and the flow rate is 0.35 mL/min throughout. Collision energies and cell accelerator voltages were optimized using sphingolipid standards with dMRM transitions as [M+H]+→[m/z = 284.3] for dihydroceramides, [M+H]+→[m/z = 287.3] for isotope-labeled dihydroceramides, [M-H2O+H]+→[m/z = 264.2] for ceramides, [MH2O+H]+→[m/z = 267.2] for isotope-labeled ceramides and [M+H]+→[M-H2O+H]+ for all targets. Sphingolipids and ceramides without available standards are identified based on HR-LC/MS, quasi-molecular ions, and characteristic product ions. Their retention times are either taken from HR-LC/MS data or inferred from the available standards. Results from LC-MS experiments are collected using Agilent Mass Hunter Workstation and analyzed using the software package Agilent Mass Hunter Quant B.07.00. Ceramide and lipid species are quantitated based on peak area ratios to the standards added to the extracts.

### Analysis of FAD and FMN by IC-HRMS

To determine the relative abundance of FAD and FMN in mouse liver tissue, extracts were prepared and analyzed by high-resolution mass spectrometry (HRMS). Approximately 20 to 30mg of tissue were pulverized in liquid nitrogen then homogenized with a Precellys Tissue Homogenizer. Metabolites were extracted using 80/20 (v/v) methanol/water with 0.1% ammonium hydroxide. Samples were centrifuged at 17,000x*g* for 5 min at 4°C, supernatants were transferred to clean tubes, followed by evaporation under vacuum. Samples were reconstituted in deionized water, then 10μL was injected into a Thermo Scientific Dionex ICS-5000+ capillary ion chromatography (IC) system containing a Thermo IonPac AS11 250×2mm 4μm column. IC flow rate was 360μL/min (at 30°C), and the gradient conditions are as follows: initial 1 mM KOH, increased to 35mM at 25 min, increased to 99mM at 39 min, and held at 99mM for 10 min. The total run time was 50min. To increase desolvation for better sensitivity, methanol was delivered by an external pump and combined with the eluent via a low dead volume mixing tee. Data were acquired using a Thermo Orbitrap Fusion Tribrid Mass Spectrometer under ESI negative mode and imported to Thermo Trace Finder software for final analysis. Relative abundance was normalized by tissue weight.

### Analysis of reduced and oxidized coenzymes by triple quadruple LC-MS

To determine the relative abundance of ubiquinone (oxidized CoQ10), ubiquinol (reduced CoQ10), ubiquinone-9 (CoQ9), and ubiquinol-9 (reduced CoQ9) in mouse liver samples, extracts were prepared and analyzed by Thermo Scientific TSQ triple quadrupole mass spectrometer coupled with a Dionex UltiMate 3000 HPLC system. Approximately 20 to 30mg of tissue were pulverized in liquid nitrogen then homogenized with a Precellys Tissue Homogenizer. Coenzymes were extracted with 500μL ice-cold 100% isopropanol. Tissue extracts were vortexed, centrifuged at 17,000x*g* for 5 min at 4°C, and supernatants were transferred to clean autosampler vials. The mobile phase was methanol containing 5mM ammonium formate. Separations of CoQ9, CoQ10, reduced-CoQ9, and reduced-CoQ10 were achieved on a Kinetex® 2.6μm C18 100 Å, 100 × 4.6mm column. The flow rate was 400μL/min at 35°C. The mass spectrometer was operated in the MRM positive ion electrospray mode with the following transitions. CoQ10/oxidized: m/z 863.7→197.1 CoQ10/reduced: m/z 882.7→197.1, CoQ9/oxidized: m/z 795.6 -> 197.1, and CoQ9/reduced: m/z 814.7 -> 197.1. Raw data files were imported to Thermo Trace Finder software for final analysis. Relative abundance was normalized by tissue weight.

### Quantification and Statistical Analysis

All measurements were taken from distinct biological samples. Unless otherwise noted, all statistical analyses were performed using GraphPad Prism (version 9) and tests described in the figure legends. In the case of multiple groups, a one- or two-way ANOVA with post-hoc tests were used to determine statistical significance. When only two groups were compared, non-parametric Mann-Whitney tests were used to determine statistical significance. Gene expression and metabolomic heatmaps were plotted as Z-scores using R(4.0.3) and ComplexHeatmap(2.6.2). The species-by-species t-test was applied for metabolomics data to identify the top differentially regulated metabolites that passed the nominal threshold p values. For multiple comparisons, the Benjamini-Hochberg procedure was used for false discovery rate (FDR) correction. Statistical analysis of energy balance was performed by ANCOVA with lean body mass as a co-variate **(Mina et al., 2018**). No statistical method was used to predetermine sample size. Unblinded analysis of histology was performed by the investigators. All data are expressed as mean ± SEM, unless otherwise specified.

## Results

### Riboflavin deficiency alters body composition and energy expenditure

In mammals, diet furnishes vitamin B2 to synthesize all the FAD for electron transfer in the mitochondria and redox reactions required for cellular homeostasis (**Powers, 2003**). Amongst key metabolic organs, *ad libitum* FAD levels were highest in the liver, heart, and kidney **(Figure 1a)**. To study how FAD depletion influences energy balance, we exposed male mice to vitamin B2-deficient (B2D) or control diets for four weeks and performed metabolic phenotyping **(Figure 1b)**. We found 99% vitamin B2 depletion (B2D) was sufficient to reduce liver FAD levels by 70% **(Figure 1c)**. Moreover, B2D significantly blunted weight gain **(Figure 1d)**, and body composition measurements showed B2D also reduced fat mass **(Figure 1e)**. When we examined the contribution of B2 to energy expenditure, we were surprised the stunted body weight phenotype of B2D did not arise from higher energy expenditure. B2D strongly reduced oxygen consumption **(Figure 1f)**, and animals moved less during nighttime measurements **(Figure 1f)**. In absolute terms, B2D-fed mice consumed the same amount of food as controls **(Figure 1f)**. Reducing B2 in the diet by 90% did not impact liver FAD levels nor influence energy balance **(Supplemental Figure S1)**. These results identify B2 requirements that make FAD available for energy expenditure requirements and body weight in male mice.

**Figure 1:**
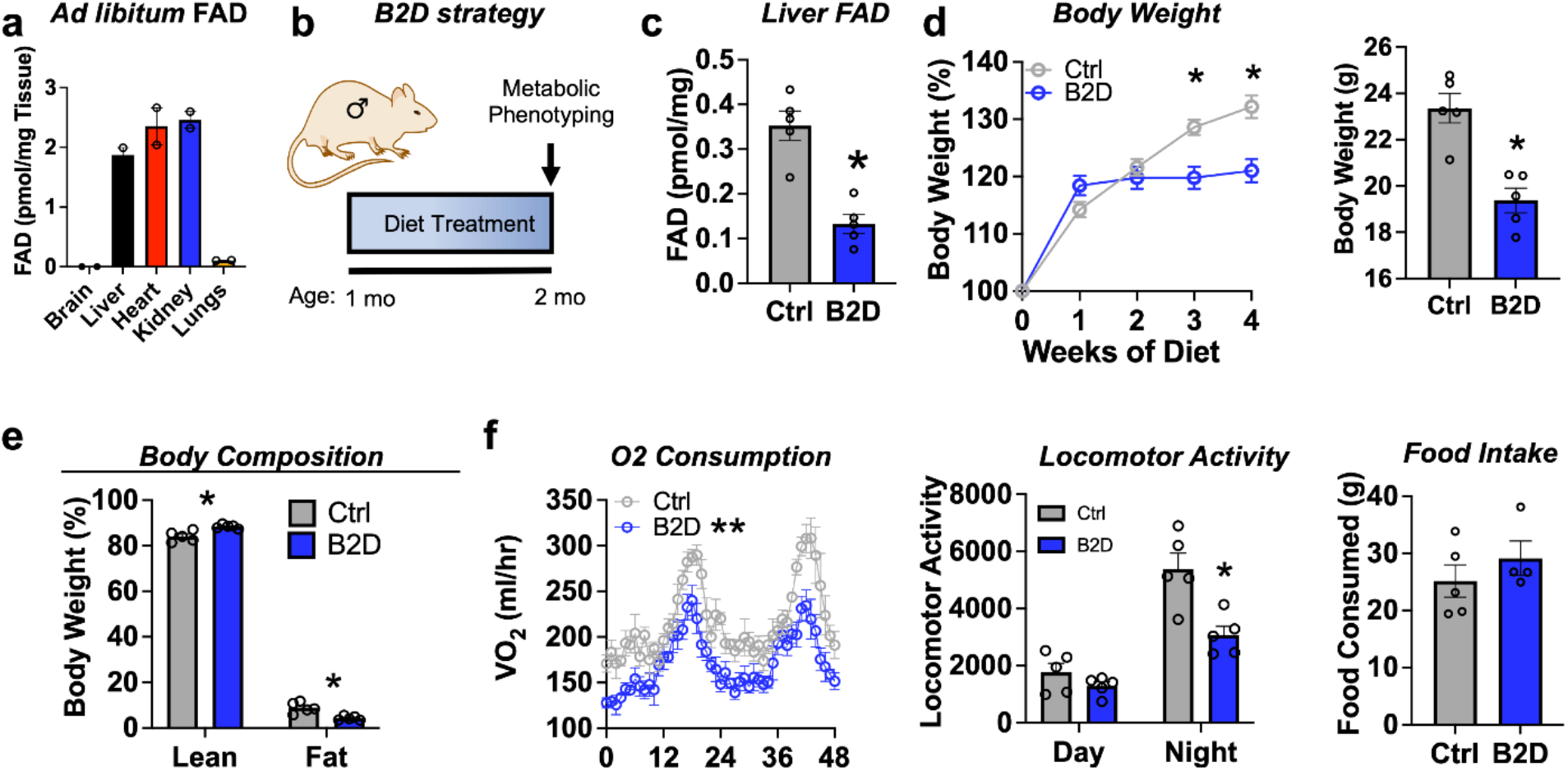
Riboflavin deficiency alters body composition and energy expenditure. **(a)** *Ad libitum* FAD concentrations measured from male WT mice. **(b)** One-month old male mice (n=4-5 per group) were exposed to 99% B2D or isocaloric control diet (Ctrl) for one month, followed by metabolic phenotyping. **(c)** FAD concentrations in the fasted liver. **(d)** Body weight (left - % initial; right – body weight at necropsy) and **(e)** body composition (% of body mass). **(f)** Mice were individually housed and monitored in CLAMS-HC. Recorded traces of oxygen consumption (VO2), locomotor activity, and cumulative food intake during time in the metabolic cages. Mann-Whitney test for liver FAD levels, body weight, and body composition **(c, d, e)**. 2-way ANOVA with Sidak multiple comparison test for body weight gain **(d)**. Statistical analysis CLAMS data was performed by ANCOVA with lean body mass as a co-variate **(f)**. Data are represented as mean +/- SEM. *p<0.05, **p<0.02.

### FAD is required for hepatic glucose production during fasting

Common clinical phenotypes of flavoprotein and nutritional riboflavin depletion disorders include fasting intolerance and hypoglycemia derived from impaired liver glucose production and metabolic flexibility (**Houten et al., 2016**). We found liver FAD displayed circadian accumulation (**Patel et al., 2012**) coinciding with the onset of gluconeogenesis (ZT12) that occurs before the active period for mice **(Figure 2a)**. We next sought to understand whether the changes in FAD that occur in the mouse liver during B2D affected gluconeogenesis. After four weeks of B2D, fasting blood glucose levels were lower **(Figure 2b)**, and further plasma analysis established higher concentrations of lactate, triglycerides (TG), free fatty acids (FFAs), and ketone bodies **(Supplemental Table S3)**. To determine the effects of B2D on gluconeogenesis *in vivo*, we subjected mice to a pyruvate tolerance test (PTT) after an overnight fast. Consistent with impaired gluconeogenesis from pyruvate, blood glucose concentrations were lower in B2D conditions compared with control diets at all times during the PTT **(Figure 2b)**. In a liver-specific way, B2D suppressed *in vivo* hepatic glucose production inferred from dilution of ^3^H-glucose infusions into fasted mice **(Figure 2b)**. These data indicate that FAD depletion directly affects liver glucose metabolism.

**Figure 2:**
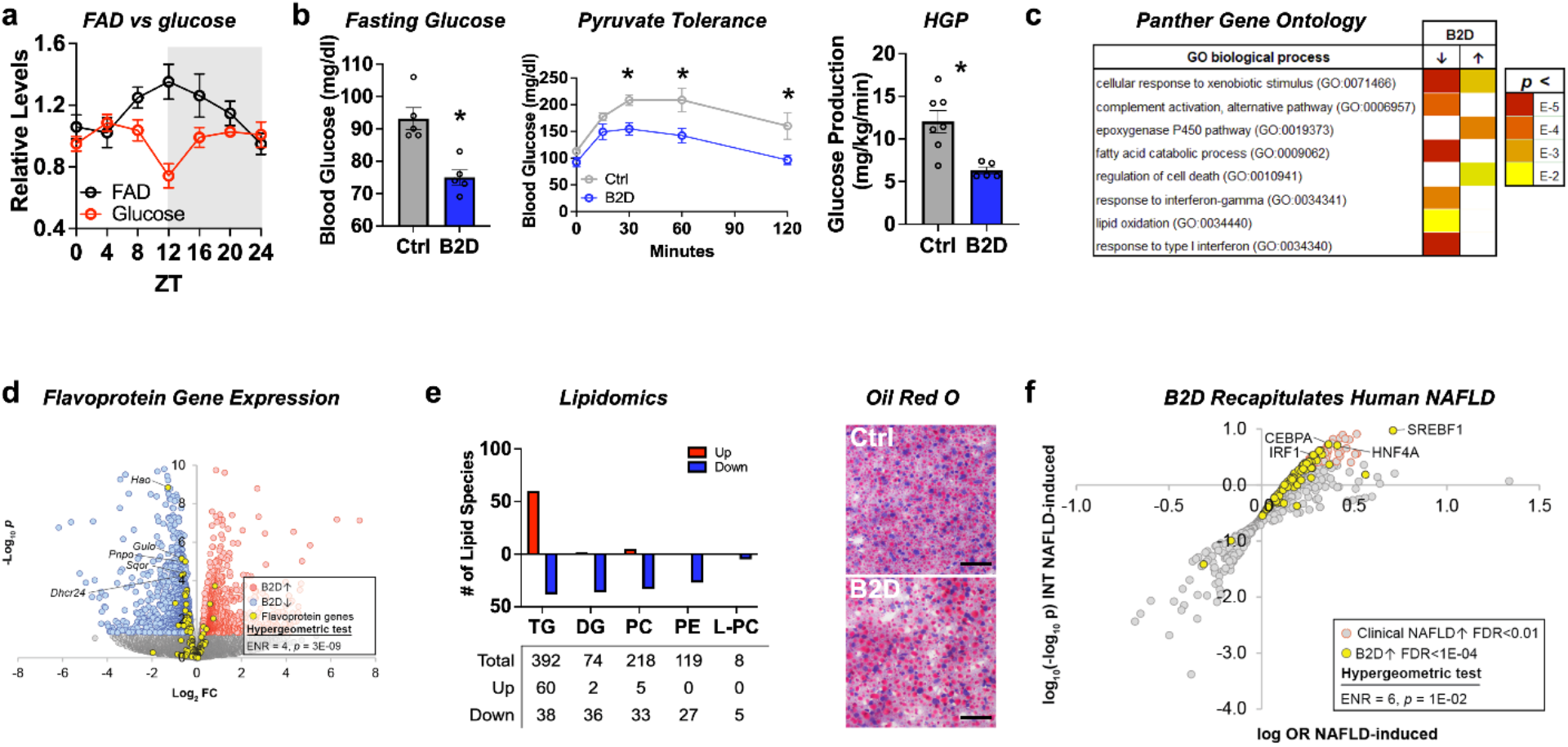
Liver glucose production requires bioavailable riboflavin. **(a)** Liver FAD and glucose levels across light/dark cycles (Zeitgeber Time, ZT) in male mice. **(b)** Overnight fasting blood glucose levels after one month of B2D or Ctrl (left) and blood glucose excursion during pyruvate tolerance tests (middle) (n=5 per group). Basal (18 h fasted) hepatic glucose production (right; n=5 B2D or n=7 Ctrl). **(c)** RNA-seq coupled with Panther Gene Ontology analysis identified pathways altered by B2D in the liver (n=5 independent animals/diet). **(d)** Volcano plot depicting expression levels of flavoprotein genes after Ctrl or B2D. **(e)** Lipidomic analysis of Ctrl and B2D (n=5 per group). This analysis identified triglycerides significantly (p<0.05) increased in B2D versus Ctrl fed mice. Representative Oil-Red-O stained liver sections from B2D or Ctrl. Scale bar 50μm. **(f)** Overlap of B2D-enriched nodes with nodes enriched in the human NAFLD gene expression consensome. The human NAFLD gene expression consensome ranks 18,162 genes based on their discovery across publicly archived clinical NAFLD case/control transcriptomic datasets. Statistical significance (*p<0.05) calculated by Mann-Whitney (**b**, left/right) or 2-way ANOVA with Sidak multiple comparison test (**b**, middle). Data are represented as mean +/- SEM.

To explore molecular outcomes of riboflavin depletion, we used RNA-seq to identify biologically cohesive gene programs of B2D in the liver. Consistent with known roles of FAD in FAO, pathways related to fatty acid catabolism and lipid oxidation were strongly repressed in response to B2D **(Figure 2c)**. Amongst the B2D-repressed genes annotated to the GO fatty acid catabolism pathway, we noted several encoded flavoproteins whose loss-of-function cause rare organic acidemias, including *Sqor* (**Friederich et al., 2020**) and *Ivd* (**Vockley and Ensenauer, 2006**). To further investigate the effect of B2D on flavoprotein gene expression, we curated a set of 117 genes encoding flavoproteins that require FAD or FMN for activity. Reflecting a specific impact of riboflavin deficiency, flavoprotein genes were enriched among B2D-repressed genes **(Figure 2d)** but not B2D-induced genes.

IEMs that arise from mutations in genes encoding mitochondria FAD transfer enzymes for FAO are identified by elevated organic acids in the blood, lipodystrophy, and fatty liver disease (**Balasubramaniam et al., 2019**). Lipidomics analysis in the liver identified phospholipid [phosphatidylethanolamine (PE), phosphatidylcholine (PC), and lyso-PC] and diacylglycerol (DG) as robustly attenuated **(Figure 2e)** in B2D-exposed mice compared to controls. Hepatic steatosis **(Figure 2e)** also accompanied B2D effects, presumably due to increased TG **(Supplemental Table S4)**. Accordingly, we hypothesized that altered expression of genes in the liver of B2D relative to normal diet controls reflected non-alcoholic fatty liver disease (NAFLD). To test this hypothesis, we performed high confidence transcriptional target (HCT) intersection analysis (**Ochsner et al., 2019**) to identify signaling nodes with significant regulatory footprints amongst liver B2D-induced or -repressed genes. From this approach, we identified strong enrichment of genes induced by B2D with metabolic transcription factor knockouts that cause fatty liver disease, such as *Ppara* (**Cotter et al., 2014; Kersten et al., 1999; Montagner et al., 2016**), *Nr1h4* (**Sinal et al., 2000**), and *Nr0b2* (**Huang et al., 2007**).

Using other computational approaches to expand upon the underpinnings of macrosteatosis caused by B2D, we retrieved nodes previously shown to contain significant transcriptional footprints within genes differentially expressed in clinical NAFLD (**Bissig-Choisat et al., 2021**). B2D-induced genes consisted of footprints for transcription factor nodes active in NAFLD **(Figure 2f)**, including HNF family members (**Xu et al., 2021**) and SREBP1 (**Shimano et al., 1997**). Collectively, our unbiased approach converged metabolic phenotypes and regulatory networks of B2D with those driving NAFLD and macrosteatosis observed in organic acidemias.

### PPARα activity maintains liver FAD pools for fasting glucose availability

Analysis of RNA-seq and ChiP-seq data (**Lee et al., 2014; Oshida et al., 2015**) discovered strongly enriched PPARα binding near promoter regions for 43 out of 121 putative flavoproteins (p=1.19×10^−11^, hypergeometric test) and supported direct coupling of FAD availability and nuclear receptor activity in the liver. Using DNA motif analysis, we also found canonical PPARα target genes among the genes downregulated by B2D **(Figure 3a)**. Likewise, we identified a set of BioGRID-curated interaction partners of PPARα that was enriched among nodes with strong footprints in B2D-repressed genes **(Figure 3a)**. Furthermore, B2D reduced PPARα–regulated flavoprotein genes **(Figure 3b)**. The observation that patterns of B2D-sensitive gene expression in the liver overlap with PPARα regulatory footprints indicated a convergent role for PPARα and riboflavin in the transcriptional control of gluconeogenic responses.

**Figure 3:**
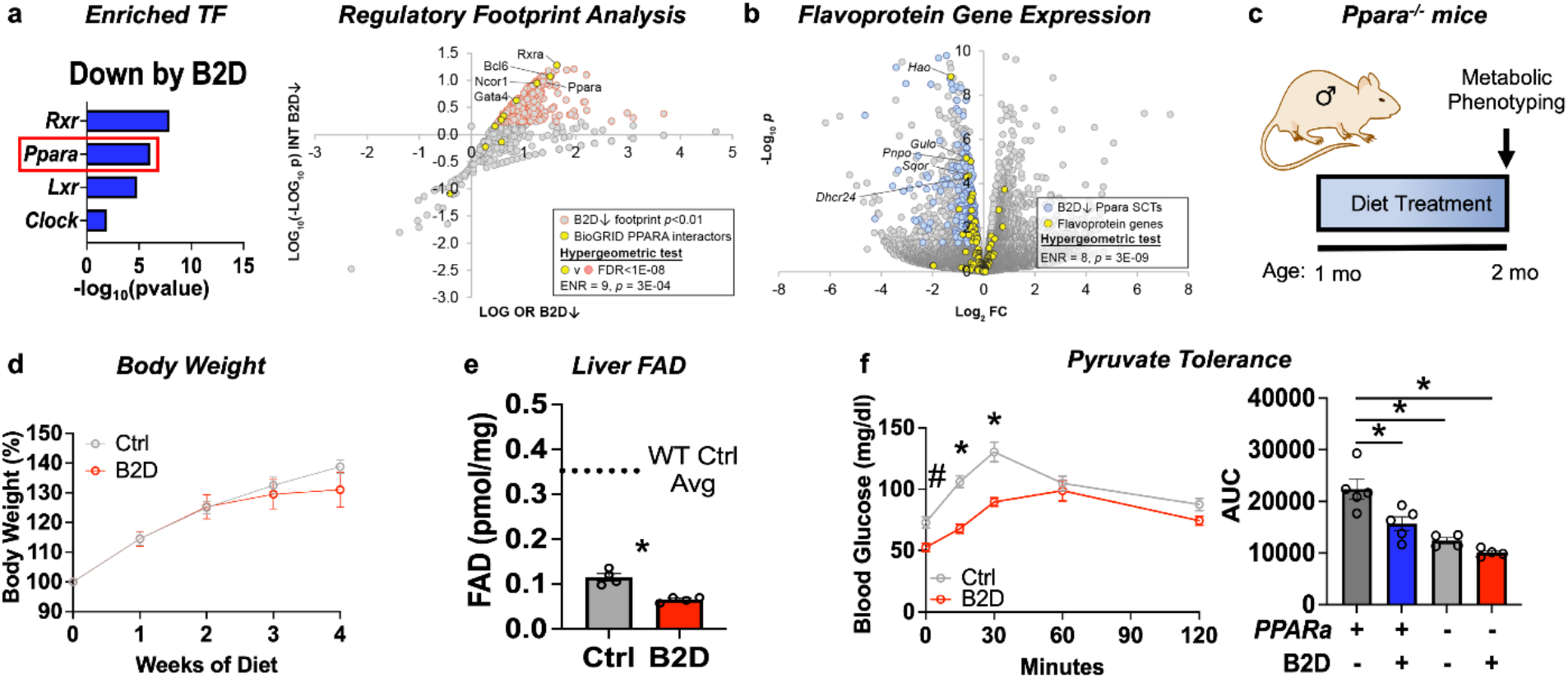
PPARα governs glucogenic responses to dietary riboflavin. **(a)** Top enriched, B2D repressed gene sets using the EnrichR transcription factor collection (left). Scatterplot showing enrichment of known BioGRID-curated PPARα interacting nodes among nodes with the most significant intersections with B2D-repressed genes (right). **(b)** Volcano plot depicting expression levels of PPARα-regulated flavoprotein genes after Ctrl or B2D. **(c)** One-month old *Ppara^-/-^* male mice (n=4 per group) were exposed to 99% B2D or isocaloric control diet (Ctrl) for one month. **(d)** Body weight (% initial) changes and **(e)** FAD concentrations in the fasted liver. FAD levels for WT Ctrl diet are shown with dashed lines. **(f)** Blood glucose levels and area under the curve (AUC) during pyruvate tolerance tests (n=4-5/group). Statistics: Mann-Whitney test **(e)**, 2-way ANOVA with Sidak multiple comparison test **(f**, left) and one-way ANOVA with Tukey’s multiple comparisons test were used to identify statistically different AUC **(f**, right). Data are represented as mean +/- SEM. *p<0.05, #p<0.10.

To explore the physiological intersections between PPARα and B2D, we phenotyped male *Ppara* whole-body knockout (pKO) mice exposed to control or B2D for one month **(Figure 3c)**. As expected (**Cotter et al., 2014**), we learned pKO largely negated effects of B2D on body weight gain **(Figure 3d)**. Baseline levels of FAD were considerably lower in pKO, and B2D exerted a more substantial suppression of FAD levels relative to historical wild-type measurements **(Figure 3e)**. PTT demonstrated B2D and pKO decreased the conversion of glucose release relative to control diets and wild-type controls **(Figure 3f)**. These findings suggest PPARα sustains FAD levels and requires riboflavin to direct glucogenic responses in the liver.

### PPARα activation by fenofibrate rescues liver glucose production even when FAD cannot be generated from diet

The FDA-approved fibrate drugs act selectively on PPARα to lower blood lipids and treat hypertriglyceridemia (**Bougarne et al., 2018**). PPARα agonists also show promise for the treatment of some mitochondrial disorders (**Steele et al., 2020**). The convergence of B2D effects on PPARα regulation of gene expression and metabolic responses suggested fenofibrate treatment may restore metabolic competency in animals on riboflavin-deficient diet. To explore this possibility, we administered fenofibrate after seven weeks of B2D exposure **(Figure 4a)**. Daily gavage with fenofibrate (300 mg/kg) for two weeks while maintaining mice on diet interventions significantly reduced body weight gain under B2D conditions **(Figure 4b)**. At the end of the experiment, fasted mice received fenofibrate two hours before blood glucose measurements. Fenofibrate increased blood glucose in both groups of mice far above pre-gavage levels **(Figure 4c)**. When liver histology was examined, we noticed hepatic steatosis was reversed by fenofibrate in B2D mice **(Figure 4d)**. Fenofibrate also decreased hepatic TG and cholesterol in both groups relative to control treatments **(Supplemental Table S4)**.

**Figure 4:**
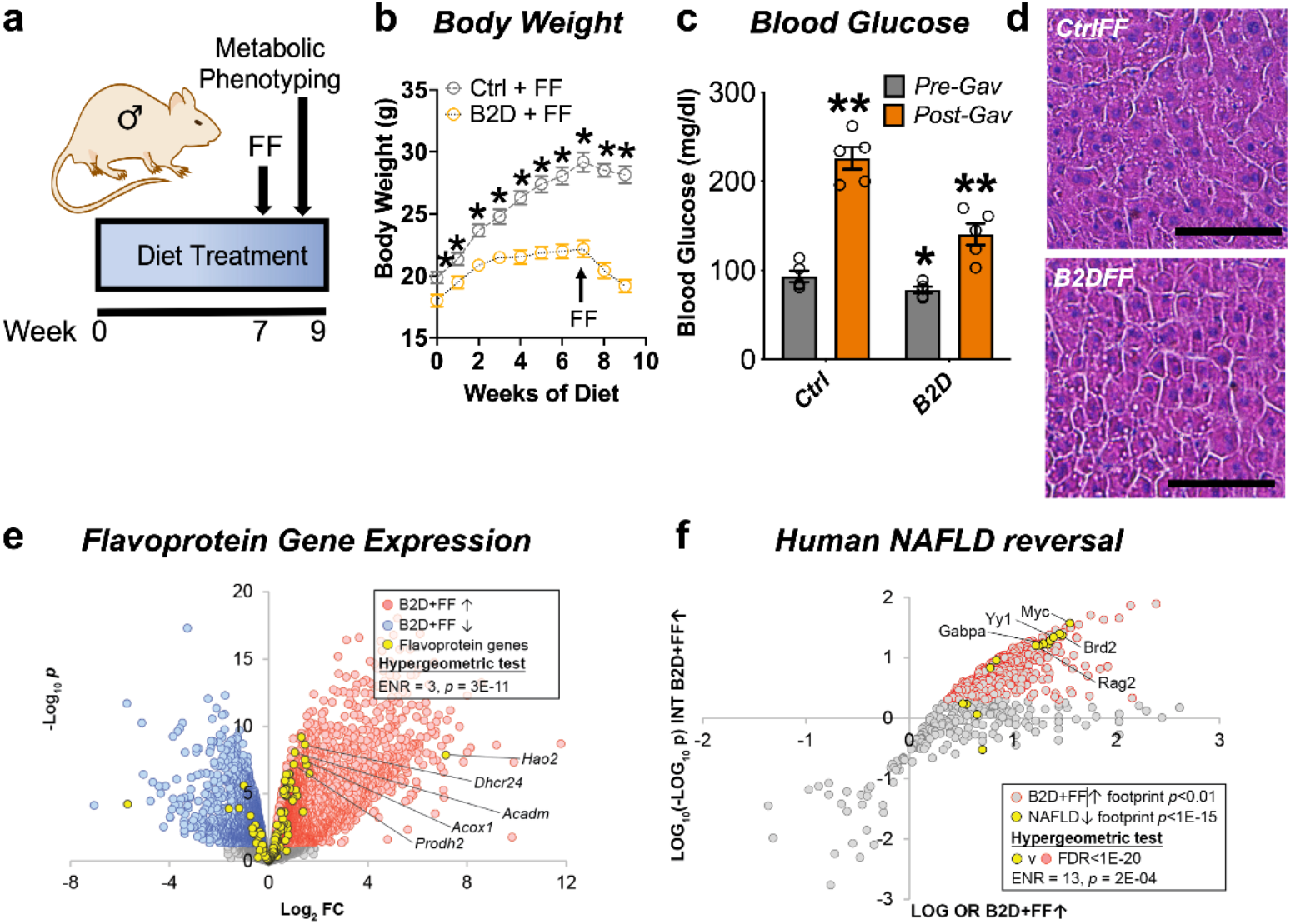
Remnant PPARα activation rescues fasting responses impaired by B2D. **(a)** One-month old male mice (n=5 per group) were exposed to 99% B2D or isocaloric control diet (Ctrl) for 9 weeks. For weeks 7-9, mice were gavaged daily with fenofibrate (FF). **(b)** Body weight (g) changes pre-gavage and during gavage. For post-gavage (week 9), mice were fasted overnight and administered FF 2 h before measurements. **(c)** Overnight fasting blood glucose levels for pre-gavage (grey) and after 2 weeks of FF treatment (orange). **(d)** Representative H&E-stained liver sections from Ctrl or B2D mice following FF treatment. Scale, 100 μm. **(e)** Expression level of flavoprotein genes in B2D-fed mice following FF. **(f)** Enrichment of B2D+FF with gene footprints repressed in human NAFLD. Statistics by two-way ANOVA corrected for multiple comparisons by Tukey **(b)**; Mann-Whitney **(c)**, Statistical enrichment shown by hypergeometric test **(e, f)**. Data are represented as mean +/- SEM. *p<0.05.

Given the ability of PPARα to regulate flavoprotein gene expression, we pursued additional RNA-seq studies to understand the mechanisms that allowed fenofibrate to rescue hypoglycemia in B2D. Consistent with restoration of the flavoprotein transcriptome in response to PPARα activation, we found the number of flavoprotein genes induced by B2D+fenofibrate more than doubled when compared to B2D alone **(Figure 4e)**. Given the improvement in hepatic steatosis after PPARα activation, we hypothesized the B2D+fenofibrate treatment caused inversion of the alignments between B2D and NAFLD transcription networks. Using this approach, we found gene footprints enriched by B2D+fenofibrate and those depleted in clinical NAFLD converged **(Figure 4f)**, including the GABP transcriptional program inactivated in inflammatory liver diseases (**Niopek et al., 2017**). These unbiased approaches strengthen the notion that PPARα activation overcomes fatty liver and hypoglycemia phenotypes imposed by B2D.

### Altered sphingolipid pools and respiratory chain efficiency in B2D

Fat and protein metabolism produces substrates for the synthesis of sphingolipids, such as ceramides and dihydroceramides, whose tissue accumulation associates with severity of fatty liver disease (**Luukkonen et al., 2016**) and mitochondrial dysfunction (**Hammerschmidt et al., 2019; Park et al., 2016**). To determine if B2D or B2D+fenofibrate also changed the sphingolipid composition of the mouse liver, we measured a battery of sphingolipids and found that deoxysphingolipids requiring alanine condensation with palmitoyl-CoA were increased by B2D and robustly decreased by fenofibrate **(Figure 5a)**. This finding may derive from the serine availability during hypoxic stress and higher NADH levels (**Yang et al., 2020**). In line with this idea, B2D conditions reduced the NAD/NADH ratio in the liver **(Figure 5b)**. Notably, deoxysphingolipids impair mitochondrial function (**Alecu et al., 2017; Muthusamy et al., 2020**) and lead to an energy deficit, especially in the liver, where fasting requires elevated demand for biomass. The observed shift in deoxysphingolipid pool size supports a critical role for fenofibrate in adapting the liver to the mitochondrial stress of B2D.

**Figure 5:**
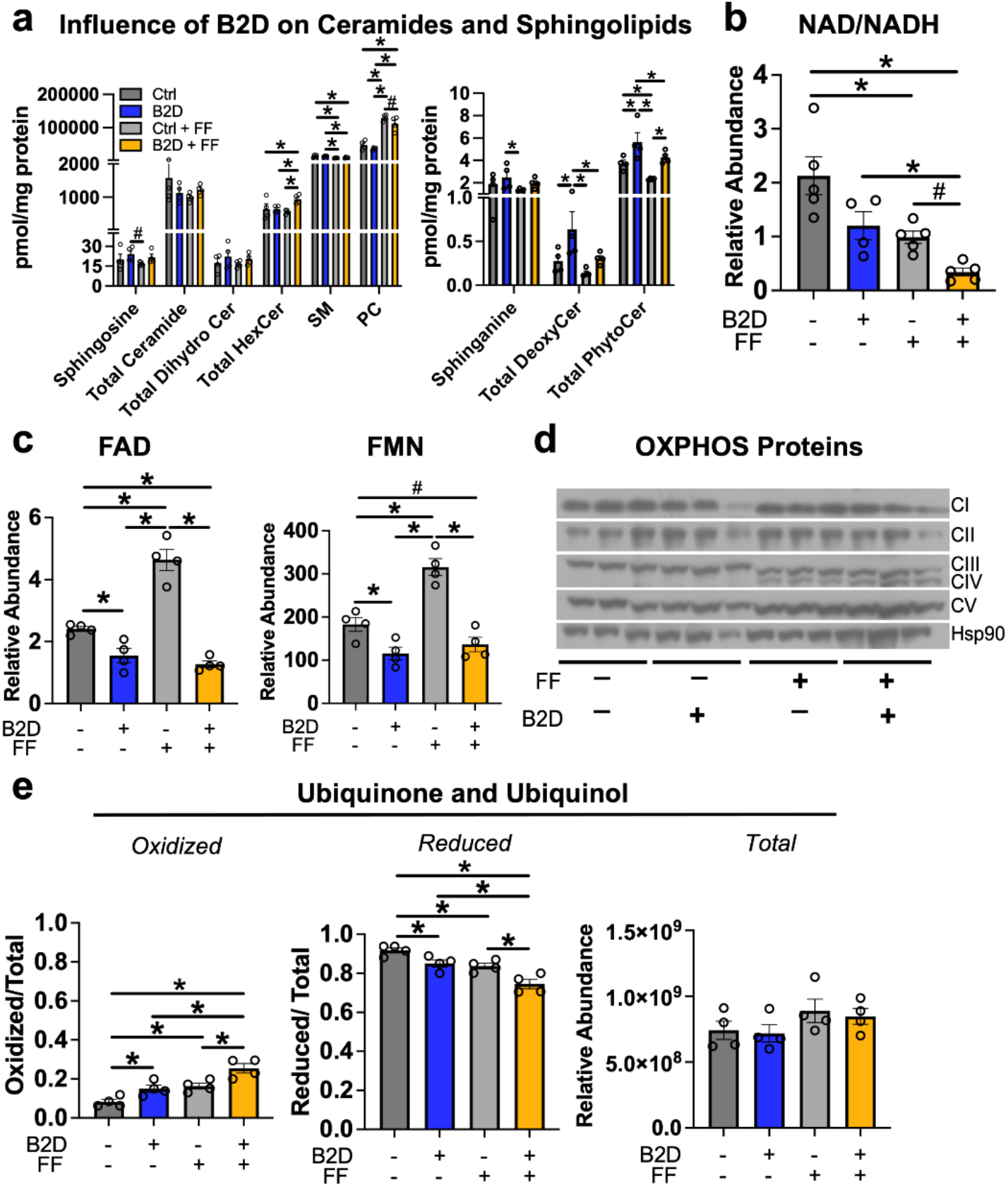
Fenofibrate impacts mitochondrial respiratory chain efficiency but does not rescue FAD levels in B2D. **(a)** Fasting liver ceramide and ceramide metabolites (pmol/mg protein) by mass spectrometry (n=4 per group). **(b)** Hepatic NAD/NADH ratio. **(c)** Fasting liver FAD and FMN relative abundance by mass spectrometry (n=4 per group). **(d)** Western blot analysis of oxidative phosphorylation complexes. Hsp90 served as the invariant control. **(e)** Ratio of oxidized and reduced liver coenzyme Q10 and Q9 by mass spectrometry (n=4 per group). *p<0.05, #p<0.1 by one-way ANOVA with post-hoc testing (Fisher’s LSD) **(a, c, e)**. Kruskal-Wallis with post-hoc testing (Dunn’s Test) **(b)**. Data are represented as mean +/- SEM.

Pathogenic variants in genes for riboflavin transport and metabolism that deplete FAD impair electron transfer from flavoenzymes in the ETC to coenzyme Q_10_ and, ultimately, the energy requirements for gluconeogenesis and fasting tolerance (**Rinaldo et al., 2002**). During prolonged fasting, beta-oxidation and proteolysis produce high fluxes of electrons that flow through the ETC. Coenzyme Q_10_ collects and converges electron flow on complex III via complex II (oxidizing succinate into fumarate) or complex I. In this way, the coenzyme Q_10_ pool accommodates variable electron fluxes and manages the mitochondria redox environment. Analysis of FAD and FMN **(Figure 5c)** confirmed fenofibrate was incapable of reconciling cofactor pools and lacked meaningful impacts on relative levels of oxidative phosphorylation proteins **(Figure 5d)**. In contrast to the reduced NAD/NADH, we found B2D treatments oxidized Coenzyme Q_10_ and rodent-biased Coenzyme Q_9_ **(Figure 5e)**, suggesting higher Q/QH2, more negative free energy for complexes I/II, and compensation for the defects in mitochondrial energy efficiency associated with B2D (**Satapati et al., 2015**).

### Fenofibrate activates the integrated stress response in B2D

We next measured concentrations of carnitines, amino acid, and hydrophilic metabolites in the liver using mass spectroscopy across B2D exposures. B2D altered the steady-state levels of valine, methionine, phenylalanine, and caused accumulation of short-chain C5 carnitines that reflect incomplete beta-oxidation of fatty acids **(Figure 6a)**. Upper arms of glucose metabolism (3PG/2PG) showed lower activities coupled with higher lactate and buildup of metabolites above pyruvate oxidation (G3P, PEP). Metaboanalyst (**Pang et al., 2021**) revealed complete removal of dietary riboflavin caused metabolite changes in the liver that enriched for organic acidemias and inborn errors of the TCA cycle **(Figure 6a)**. Moreover, B2D caused accumulation of oxidized TCA cycle metabolic intermediates, including fumarate, malate, and 2HG. These findings were compatible with gene sets **(Figure 6b)** associated with hypoxia and epithelial-mesenchymal transition (**Sciacovelli et al., 2016; Ward et al., 2010**).

**Figure 6:**
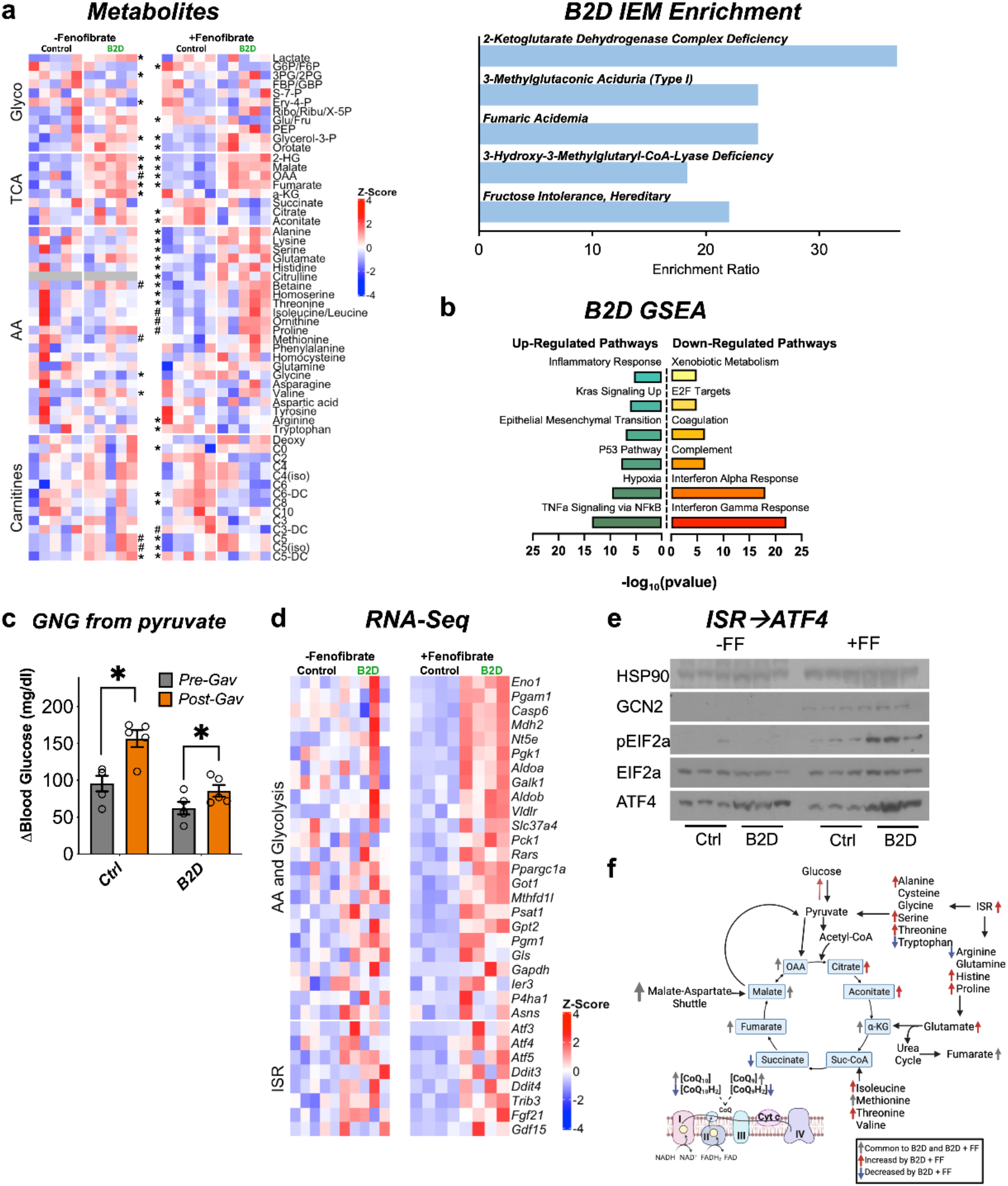
ISR activity forms the basis to reconcile FAD disruption. **(a)** Glycolysis, TCA, amino acid, fatty acid oxidation (carnitines) metabolites measured by mass spectrometry across B2D and FF treatments (left; shown as Z-score). Metaboanalyst integration demonstrates B2D causes organic acidemias (top right). **(b)** RNA-seq coupled with gene set enrichment analysis identified gene signatures altered by B2D in the liver (n = 5 independent animals/diet). **(c)** Change in glucose levels 30 minutes after pyruvate injection pre- and post-fenofibrate gavage (n=5). **(d)** Amino acid, glucose metabolism, and integrated stress response (ISR) genes in B2D or B2D + FF, shown as Z-score from RNA-seq data. **(e)** Western blot analysis for ISR proteins in liver lysates from mice treated with B2D or B2D+FF and fasted overnight. **(f)** B2D + FF activates the ISR, increasing amino acids that restore glucose availability. Statistics by two-way ANOVA and post-hoc tests (Fisher LSD) **(c)**; Mann-Whitney **(a, d)**. Data are represented as mean +/- SEM. *p<0.05, #p<0.10.

PPARα activity favors conversion of proteins to provide amino acids as substrates for anabolic processes (**Kersten et al., 2001**). Our phenotyping analysis of fenofibrate suggested alternative carbon sources might supply the substrates to support glucose production during B2D **(Figure 4c)**, including pyruvate, which recovered blood glucose in B2D to pre-gavage control levels **(Figure 6c)**. Amino acids, such as alanine and serine, are also significant contributors to de novo synthesis of glucose. In line with this idea, gluconeogenic amino acids (serine, glutamate, histidine, alanine) showed selective and unique accumulation in the combined B2D and fenofibrate treatments (**Figure 6a**). Steady-state levels of other anaplerotic amino acids that replenish the TCA cycle, isoleucine, and threonine, also increased at the expense of reduced ketosis and β-hydroxybutyrate **(Figure 6a** and **Supplemental Table S3)**. Similarly, B2D+fenofibrate elevated levels of TCA cycle metabolites citrate and aconitate and increased the fumarate to succinate ratio. Importantly, we noted moderated levels of carnitines enriched in organic acidemias (C5 and C6). These conditions suggest PPARα activation inhibits cataplerosis of TCA cycle intermediates.

The constellation of phenotypic effects resulting from riboflavin depletion and fenofibrate suggested global shifts in gene expression and metabolism. RNA-seq indicated B2D+fenofibrate precipitated a response with elements of greater amino acid and glucose metabolism **(Figure 6d)**. We also observed a gene signature for the integrated stress response (ISR) further supported by increased expression of the master transcription factor regulator *Atf4*, as well as its key target genes (*Psat1, Fgf21, Asns, Gpt2, Ddit3*) during combined B2D+fenofibrate treatments. Mechanistically, ISR activation occurs through phosphorylation of eIF2α and ATF4 translation, which mediates transcription of target genes to resolve the ISR and regain amino acid homeostasis (**Harding et al., 2003; Ye et al., 2012**). In line with this idea, B2D+fenofibrate achieved robust GCN2 expression, eIF2α phosphorylation, and higher ATF4 levels relative to any other treatment **(Figure 6e)**. Together, our integrated transcription and mass spectrometry analyses favor a model in which PPARα activation precipitates concerted activation of the ISR, which in turn overcomes the fasting intolerance imposed by B2D **(Figure 6f)**.

## Discussion

Our work sheds light on how the liver copes with severe metabolic crises and the FAO disorders caused by flavoprotein disruption or FAD depletion during prolonged fasting when both beta-oxidation and gluconeogenesis are concomitantly activated. We speculate these FAO disorders rely on conservation responses to produce energy when the mitochondria are starved of FAD and FMN required for ETC activity. Interestingly, the metabolic phenotype of B2D resists conditions unfavorable for mitochondrial function, including hypoxia, suggesting that these changes select for survival. Increased reliance upon glycolysis occurs in mitochondrial disease (**Robinson, 2006**) and hypoxia responses activate glycolytic enzymes that allow energy production when the mitochondria are starved of oxygen as a substrate for oxidative phosphorylation. Likewise, our data support the idea that the stress of B2D gives rise to an environment for PPARα to co-opt the ISR, adapt to TCA cycle dysfunction, and engage hypoxia enzymes to reconcile anaplerosis. Elevated TCA cycle intermediates, such as fumarate, then act to stabilize antioxidant transcription factors and protect against oxidative stress in the liver (**Ashrafian et al., 2012**) during the stress of IEMs modeled in our study.

Mammals cannot synthesize vitamin B2, so diet remains the only available source of FAD and FMN (**Powers, 2003**). While the effect of B2D may act through different pathways and tissues, we demonstrated some of these effects may be mediated through disturbance of nuclear receptor activity and altered glucose availability. One important caveat of these experiments is that the kidney also contributes to glucose production during fasting (**Joseph et al., 2000**). While our *in-vivo* experiments do not explore whether the kidney reconciled any liver gluconeogenic deficiency, our observations reveal fundamental vitamin requirements to source the liver with the FAD and FMN pools necessary for energy balance.

The energetic requirements of fasting dictate substrate oxidation and electron transport. B2D restricts mitochondrial function, which causes reductive pressures that exacerbate ROS formation in part through proton leak. PPARα activation prevents oxidative stress (**Ip et al., 2004**) by merging lower NAD/NADH and oxidized Q, which, in turn, lowers the free energy of electron transport through complexes I and II. These data are consistent with inhibition of complex II, resulting in a more oxidized Q pool and reveals an important adaptation that allows the liver to overcome riboflavin deficiency (**Treberg et al., 2011**).

IEMs modeled in our study frequently present NAFLD-like phenotypes that contribute to fasting intolerance (**Rinaldo et al., 2002**). In our model, B2D alters lipid profiles and gene expression patterns that converge B2D with more common NAFLD phenotypes. The liver dominates mass-specific metabolic rates (**Rolfe and Brown, 1997**) and, for this reason, the NAFLD caused by B2D likely reflects a combination of lower hepatic fatty acid oxidation contributing to the accumulation of liver triglycerides and other complex lipid species observed in obesity (**Moore et al., 2022**). Our lipidomics also revealed that B2D caused accumulation of deoxysphingolipids, which also become more abundant in NAFLD from incomplete fat oxidation and accrual of toxic intermediates (**Gai et al., 2020**). These findings are particularly relevant for discovering new biomarkers for fatty liver disease.

Inefficient amino acid availability, or increased requirements of amino acids to maintain gluconeogenesis, activate ATF4 and the ISR. ATF4 is the principal downstream effector of the ISR, whose regulation becomes altered in human and rodent NAFLD (**Puri et al., 2008; Seo et al., 2009**). Integrated metabolomic and RNA-seq studies demonstrated that fenofibrate unveils the ISR to increase the abundance of anaplerotic amino acids. We are unaware of other studies that observe coincident activation of PPARα and ATF4 to drive adaptive responses to liver stress. However, induction of shared PPARα and ATF4 targets, including *Fgf21*, may contribute to hepatoprotection from lipotoxic lipid accumulation (**Montagner et al., 2016**). It will be interesting in the future to determine why B2D exposes a vulnerability to fenofibrate that engages selective genome regulation by PPARα and ATF4 for recovering glucose production in settings of fasting intolerance.

This study is unique in its comprehensive approach to understand the complex metabolic consequences of FAD depletion and riboflavin auxotrophy in mouse models. While further studies are needed, we describe allostatic outcomes of PPARα activation that overcome bioenergetic costs of fasting. Rare diseases of flavoprotein mutations, including organic acidemias, cause substantial morbidity and have no cure. Therefore, understanding how nuclear receptor regulation of flavoprotein function and FAD pool distribution surmounts hypoglycemia and fatty liver is valuable for implementing metabolic interventions and future therapeutic strategies.

## Acknowledgements

This work was funded by the Nancy Chang, Ph.D. Award for Research Excellence at Baylor College of Medicine, American Diabetes Association #1-18-IBS-105 and NIH R01DK114356 (to SMH).

## Author contributions

PMM and SMH conceptualized the study. PMM, ARC, PKS, and SMH designed experiments. PMM, NJM, and SMH wrote the manuscript with editorial input from all authors. PMM performed all experiments with assistance as noted: PKS, KHK, RS, JBF, XL, ANA, and ARC assisted with mouse phenotyping; JHP and BK helped interpret ETC function; VP, NP, LT, PLL, LW, and SAS provided metabolomics analysis and support; SO and NJM assisted with RNA-seq analysis and data integration. S.M.H. is the guarantor of this work and, as such, had full access to all the data in the study and takes responsibility for the integrity of the data.

## Materials Availability

This study did not generate new unique reagents. The authors declare that reagents utilized are available upon reasonable request to the corresponding author.

## Data Availability

All data generated or analyzed during this study are included in the published article and supplemental files. RNA-sequencing datasets generated have been deposited into the National Center for Biotechnology Gene Expression Omnibus database. The accession numbers for these datasets are NCBI GEO: GSE206200.

## Supplemental Materials

### Supplemental Figure

**Supplemental Figure S1, Related to Figure 1.**
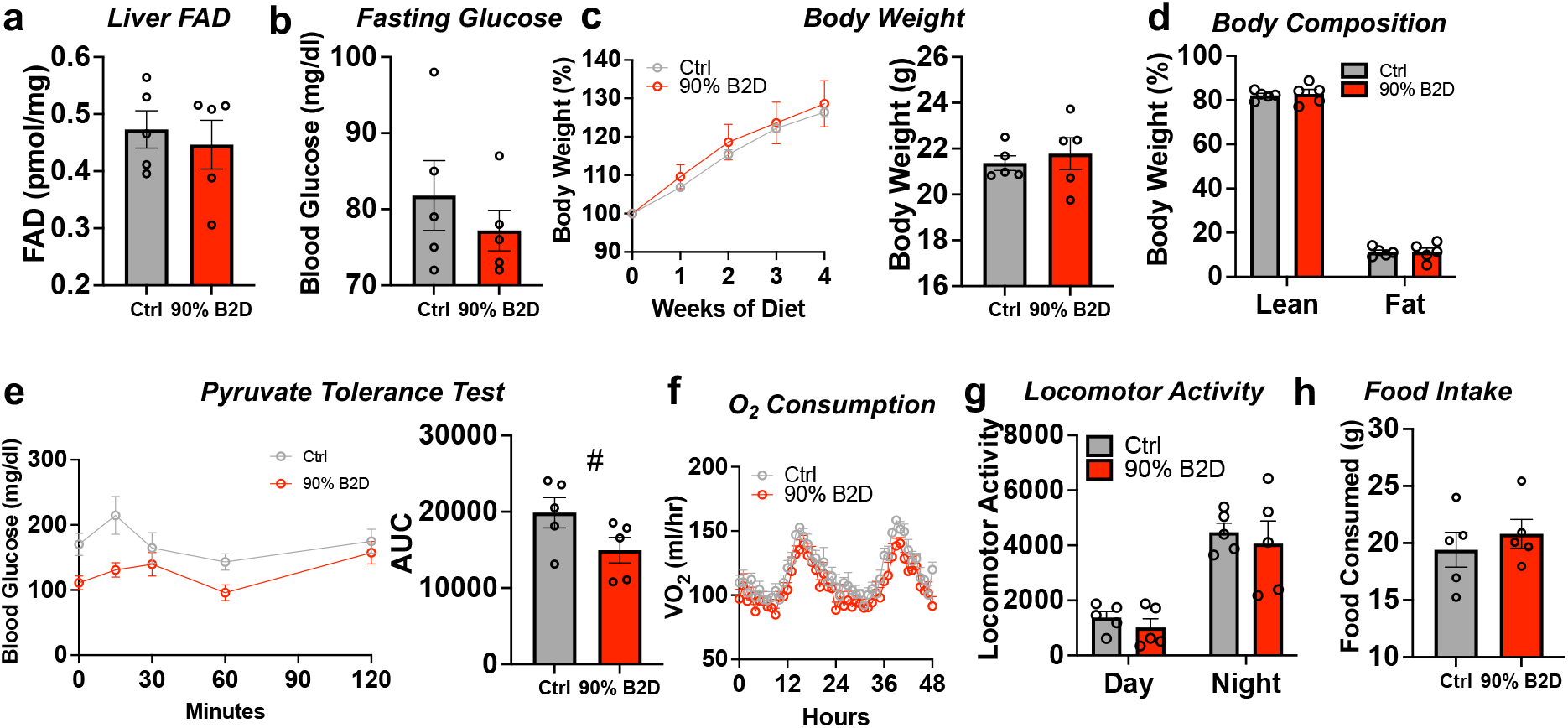
Metabolic effects of 90% riboflavin deficiency in mice. **(a)** FAD concentrations in the fasted liver. **(b)** Overnight fasting blood glucose levels after one month on 90% B2D or Ctrl. **(c)** Body weight changes and **(d)** body composition (as % of body mass). **(e)** Blood glucose excursion in 90% B2D or Ctrl mice during pyruvate tolerance tests (n=5). Statistical significance for AUC by Mann-Whitney test, #p<0.10. **(f)** Mice were individually housed and monitored in CLAMS home cages. Recorded traces of oxygen consumption (VO2, ml/hl). **(g)** Locomotor activity and **(h)** cumulative food intake (g) during time in metabolic cages. Data represented as mean +/- SEM.

### Supplemental Tables

**Supplemental Table S1.** Macronutrient compositions of 90% B2D used in the study.

|  | **90% Ctrl** | **90% Deficient** |
| --- | --- | --- |
| Research Diets Name | AIN-93G Diet | AIN-93G Diet With No Added Riboflavin |
| Research Diets Number | D10012G | D12030102 |
| **Riboflavin Concentration** | **0.00698 g/kg** | **0.00098 g/kg** |
| **Formula** | **g/kg** | **g/kg** |
| Casein (0.00049g/kg riboflavin) | 200 | 200 |
| L-Cystine | 3 | 3 |
| Sucrose | 100 | 100 |
| Corn Starch | 397.486 | 397.486 |
| Maltodextrin 10 | 132 | 132 |
| Cellulose | 50 | 50 |
| Soybean Oil | 70 | 70 |
| t-butylrhydroquinone | 0.014 | 0.014 |
| Choline Bitartrate | 2.5 | 2.5 |
| Mineral Mix S10022G | 35 | 35 |
| Vitamin Mix V10037 (0.006 g/kg Riboflavin) | 10 | 0 |
| Vitamin Mix V15920, No Added Riboflavin | 0 | 10 |
| **% kcal from** |  |  |
| Protein | 20.3 | 20.3 |
| Carbohydrates | 63.9 | 63.9 |
| Fat | 7 | 7 |
| Kcal/g | 4 | 4 |

**Supplemental Table S2.** Macronutrient compositions of 99%B2D used in the study.

|  | **99% Ctrl** | **99% Deficient (B2D)** |
| --- | --- | --- |
| Research Diets Name | Modified AIN-93G Diet<br>With L-Amino Acids | Modified AIN-93G Diet With L-Amino<br>Acids and No Added Riboflavin |
| Research Diets Number | A18041301 | A19080901 |
| **Riboflavin Concentration** | **0.006 g/kg** | **0 g/kg** |
| **Formula** | **g/kg** | **g/kg** |
| L-Alanine | 5 | 5 |
| L-Arginine | 5.9 | 5.9 |
| L-Asparagine, monohydrate | 7 | 7 |
| L-Aspartic Acid | 5 | 5 |
| L-Cysteine | 4.2 | 4.2 |
| L-Glutamic Acid | 20.7 | 20.7 |
| L-Glutamine | 17.1 | 17.1 |
| Glycine | 3 | 3 |
| L-Histidine-HCl, monohydrate | 4.5 | 4.5 |
| L-Isoleucine | 7.5 | 7.5 |
| L-Leucine | 15.7 | 15.7 |
| L-Lysine, HCl | 13.1 | 13.1 |
| L-Methionine | 5 | 5 |
| L-Phenylalanine | 8.4 | 8.4 |
| L-Proline | 17.6 | 17.6 |
| L-Serine | 9.9 | 9.9 |
| L-Threonine | 7.1 | 7.1 |
| L-Tryptophan | 2.1 | 2.1 |
| L-Tyrosine | 9.1 | 9.1 |
| L-Valine | 9.2 | 9.2 |
| Sucrose | 107.0777 | 107.0777 |
| Corn Starch | 397.486 | 397.486 |
| Maltodextrin 10 | 132 | 132 |
| Cellulose | 50 | 50 |
| Soybean Oil | 70 | 70 |
| t-butylrhydroquinone | 0.014 | 0.014 |
| Calcium Carbonate | 7.34 | 7.34 |
| Potassium Citrate 1, monohydrate | 2.4773 | 2.4773 |
| Potassium Phosphate, monobasic | 6.86 | 6.86 |
| Calcium Phosphate, dibasic | 7 | 7 |
| Sodium Chloride | 2.59 | 2.59 |
| Sodium Bicarbonate | 7.5 | 7.5 |
| Choline Bitartrate | 2.5 | 2.5 |
| Mineral Mix S10022C | 3.5 | 3.5 |
| Vitamin Mix V10037 (0.006 g/kg Riboflavin) | 10 | 0 |
| Vitamin Mix V15920, No added riboflavin | 0 | 10 |
| **% kcal from** |  |  |
| Protein | 18 | 18 |
| Carbohydrates | 65.9 | 65.9 |
| Fat | 7.1 | 7.1 |
| Kcal/g | 3.924 | 3.924 |

**Supplemental Table S3, Related to Figure 2.** Serum parameters in B2D interventions. Data are mean ± SEM. The number of animals in each group are indicated in parentheses. Statistical significance (p) was determined by Mann-Whitney tests.

|  | **Cholesterol** | **TGs** | **Lactate** | **FFA** | **b-HB** | **Glycerol** |
| --- | --- | --- | --- | --- | --- | --- |
|  | **(mg/dl)** | **(mg/dl)** | **(mM)** | **(μM)** | **(mM)** | **(μM)** |
| 90% Ctrl | 122.1 ± 6.4 (5) | 152.4 ± 17.8 (5) | 5.2 ± 0.3 (5) | NA | NA | NA |
| 90% B2D | 120.0 ± 8.4 (4) | 113.6 ± 25.5 (5) | 3.5 ± 0.3 (5) | | | |
| p | 0.84 | 0.25 | 0.25 |  |  |  |
| 99% Ctrl | 146.5 ± 7.2 (5) | 100.9 ± 6.7 (5) | 4.5 ± 0.5 (5) | 994.1 ± 84.9 (5) | 0.19 ± 0.02 (5) | 105.4 ± 15.4 (5) |
| 99% B2D | 134.3 ± 11.0 (4) | 126.6 ± 17.7 (4) | 6.4 ± 0.4 (4) | 1128.5 ± 39.0 (5) | 0.27 ± 0.03 (5) | 134.0 ± 13.3 (4) |
| p | 0.36 | 0.18 | 0.18 | 0.15 | 0.08 | 0.22 |
| pKO Ctrl | 172.8 ± 15.0 (4) | 120.7 ± 15.5 (4) | 2.4 ± 0.1 (4) | 3092.9 ± 501 (4) | 0.06 ± 0.01 (4) | 157.9 ± 10.5 (4) |
| pKO B2D | 142.9 ± 10.7 (4) | 212.9 ± 31.8 (4) | 1.9 ± 0.1 (4) | 2660.9 ± 144 (4) | 0.08 ± 0.04 (4) | 131.1 ± 8.4 (3) |
| p | 0.16 | **0.04** | **0.04** | 1.00 | 0.52 | 0.12 |
| Ctrl + FF | 181.6 ± 16.5 (5) | 40.7 ± 3.9 (5) | 3.3 ± 0.5 (5) | 1136.3 ± 120 (5) | 0.22 ± 0.02 (5) | 99.3 ± 10.4 (5) |
| B2D + FF | 194.1 ± 26.7 (5) | 39.4 ± 3.8 (5) | 3.7 ± 0.5 (4) | 1142.6 ± 118 (5) | 0.13 ± 0.03 (5) | 76.6 ± 10.6 (5) |
| p | 0.70 | 0.82 | 0.82 | 0.84 | **0.04** | 0.16 |

**Supplemental Table S4, Related to Figure 4.** Liver triglycerides and cholesterol in B2D interventions. Data are mean ± SEM. The number of animals in each group are indicated in parentheses. Statistical significance (p) was determined Mann-Whitney tests.

|  | <u>TGs (mg/g)</u> |  | <u>Cholesterol (mg/g)</u> |  |
| --- | --- | --- | --- | --- |
|  |  | **p** |  | **p** |
| 99% Ctrl | 22.4 ± 2.1 (4) | **0.05** | 1.1 ± 0.2 (4) | 0.26 |
| 99% B2D | 32.08 ± 3.45 (3) | | 0.8 ± 0.1 (4) | |
| Ctrl + FF | 8.3 ± 1.4 (4) | 0.24 | 0.7 ± 0.1 (4) | **0.02** |
| B2D + FF | 11.8 ± 2.3 (4) | | 0.2 ± 0.1 (4) | |

**Supplemental Table S5:** Antibodies used in this study.

| Antibody | Company | Catalog Number | RRID |
| --- | --- | --- | --- |
| Hsp90 | Cell Signaling | C45G5 | AB_2233307 |
| Gcn2 | Cell Signaling | 3302 | AB_2277617 |
| p-Eif2a | Cell Signaling | 9721 | AB_330951 |
| Eif2a | Santa Cruz Biotech | Sc133132 | AB_1562699 |
| Atf4 | Cell Signaling | 11815 | AB_2616025 |
| Total OXPHOS<br>Rodent Cocktail | Abcam | Ab110413 | AB_2629281 |
| Rabbit HRP | Cell Signaling | 7074S | AB_2099233 |
| Mouse HRP | Cell Signaling | 7076P2 | AB_330924 |

## References

Alecu, I., Tedeschi, A., Behler, N., Wunderling, K., Lamberz, C., Lauterbach, M.A., Gaebler, A., Ernst, D., Van Veldhoven, P.P., Al-Amoudi, A., et al. (2017). Localization of 1-deoxysphingolipids to mitochondria induces mitochondrial dysfunction. J Lipid Res 58, 42–59. 10.1194/jlr.M068676

Ashrafian, H., Czibik, G., Bellahcene, M., Aksentijević, D., Smith, A.C., Mitchell, S.J., Dodd, M.S., Kirwan, J., Byrne, J.J., Ludwig, C., et al. (2012). Fumarate is cardioprotective via activation of the Nrf2 antioxidant pathway. Cell Metab 15, 361–371. 10.1016/j.cmet.2012.01.017

Balasubramaniam, S., Christodoulou, J., and Rahman, S. (2019). Disorders of riboflavin metabolism. J Inherit Metab Dis 42, 608–619. 10.1002/jimd.12058

Bissig-Choisat, B., Alves-Bezerra, M., Zorman, B., Ochsner, S.A., Barzi, M., Legras, X., Yang, D., Borowiak, M., Dean, A.M., York, R.B., et al. (2021). A human liver chimeric mouse model for non-alcoholic fatty liver disease. JHEP Rep 3, 100281. 10.1016/j.jhepr.2021.100281

Bougarne, N., Weyers, B., Desmet, S.J., Deckers, J., Ray, D.W., Staels, B., and De Bosscher, K. (2018). Molecular actions of PPARα in lipid metabolism and inflammation. Endocr Rev 39, 760–802. 10.1210/er.2018-00064

Chaurasia, B., Tippetts, T.S., Mayoral Monibas, R., Liu, J., Li, Y., Wang, L., Wilkerson, J.L., Sweeney, C.R., Pereira, R.F., Sumida, D.H., et al. (2019). Targeting a ceramide double bond improves insulin resistance and hepatic steatosis. Science 365, 386–392. 10.1126/science.aav3722

Cotter, D.G., Ercal, B., d’Avignon, D.A., Dietzen, D.J., and Crawford, P.A. (2014). Impairments of hepatic gluconeogenesis and ketogenesis in PPARα-deficient neonatal mice. Am J Physiol Endocrinol Metab 307, E176–185. 10.1152/ajpendo.00087.2014

Friederich, M.W., Elias, A.F., Kuster, A., Laugwitz, L., Larson, A.A., Landry, A.P., Ellwood-Digel, L., Mirsky, D.M., Dimmock, D., Haven, J., et al. (2020). Pathogenic variants in SQOR encoding sulfide:quinone oxidoreductase are a potentially treatable cause of Leigh disease. J Inherit Metab Dis 43, 1024–1036. 10.1002/jimd.12232

Gai, Z., Gui, T., Alecu, I., Lone, M.A., Hornemann, T., Chen, Q., Visentin, M., Hiller, C., Hausler, S., and Kullak-Ublick, G.A. (2020). Farnesoid X receptor activation induces the degradation of hepatotoxic 1-deoxysphingolipids in non-alcoholic fatty liver disease. Liver Int 40, 844–859. 10.1111/liv.14340

Hammerschmidt, P., Ostkotte, D., Nolte, H., Gerl, M.J., Jais, A., Brunner, H.L., Sprenger, H.G., Awazawa, M., Nicholls, H.T., Turpin-Nolan, S.M., et al. (2019). CerS6-derived sphingolipids interact with Mff and promote mitochondrial fragmentation in obesity. Cell 177, 1536–1552.e1523. 10.1016/j.cell.2019.05.008

Harding, H.P., Zhang, Y., Zeng, H., Novoa, I., Lu, P.D., Calfon, M., Sadri, N., Yun, C., Popko, B., Paules, R., et al. (2003). An integrated stress response regulates amino acid metabolism and resistance to oxidative stress. Mol Cell 11, 619–633. 10.1016/s1097-2765(03)00105-9

Houten, S.M., Violante, S., Ventura, F.V., and Wanders, R.J. (2016). The biochemistry and physiology of mitochondrial fatty acid β-oxidation and its genetic disorders. Annu Rev Physiol 78, 23–44. 10.1146/annurev-physiol-021115-105045

Huang, J., Iqbal, J., Saha, P.K., Liu, J., Chan, L., Hussain, M.M., Moore, D.D., and Wang, L. (2007). Molecular characterization of the role of orphan receptor small heterodimer partner in development of fatty liver. Hepatology 46, 147–157. 10.1002/hep.21632

Ip, E., Farrell, G., Hall, P., Robertson, G., and Leclercq, I. (2004). Administration of the potent PPARalpha agonist, Wy-14,643, reverses nutritional fibrosis and steatohepatitis in mice. Hepatology 39, 1286–1296. 10.1002/hep.20170

Jackevicius, C.A., Tu, J.V., Ross, J.S., Ko, D.T., Carreon, D., and Krumholz, H.M. (2011). Use of fibrates in the United States and Canada. Jama 305, 1217–1224. 10.1001/jama.2011.353

Joseph, S.E., Heaton, N., Potter, D., Pernet, A., Umpleby, M.A., and Amiel, S.A. (2000). Renal glucose production compensates for the liver during the anhepatic phase of liver transplantation. Diabetes 49, 450–456. 10.2337/diabetes.49.3.450

Kersten, S., Mandard, S., Escher, P., Gonzalez, F.J., Tafuri, S., Desvergne, B., and Wahli, W. (2001). The peroxisome proliferator-activated receptor alpha regulates amino acid metabolism. Faseb j 15, 1971–1978. 10.1096/fj.01-0147com

Kersten, S., Seydoux, J., Peters, J.M., Gonzalez, F.J., Desvergne, B., and Wahli, W. (1999). Peroxisome proliferator-activated receptor alpha mediates the adaptive response to fasting. J Clin Invest 103, 1489–1498. 10.1172/jci6223

Kettner, N.M., Voicu, H., Finegold, M.J., Coarfa, C., Sreekumar, A., Putluri, N., Katchy, C.A., Lee, C., Moore, D.D., and Fu, L. (2016). Circadian homeostasis of liver metabolism suppresses hepatocarcinogenesis. Cancer Cell 30, 909–924. 10.1016/j.ccell.2016.10.007

Kim, K.H., Choi, J.M., Li, F., Dong, B., Wooton-Kee, C.R., Arizpe, A., Anakk, S., Jung, S.Y., Hartig, S.M., and Moore, D.D. (2019). Constitutive androstane receptor differentially regulates bile acid homeostasis in mouse models of intrahepatic cholestasis. Hepatol Commun 3, 147–159. 10.1002/hep4.1274

Lee, J.M., Wagner, M., Xiao, R., Kim, K.H., Feng, D., Lazar, M.A., and Moore, D.D. (2014). Nutrient-sensing nuclear receptors coordinate autophagy. Nature 516, 112–115. 10.1038/nature13961

Lee, S.S., Pineau, T., Drago, J., Lee, E.J., Owens, J.W., Kroetz, D.L., Fernandez-Salguero, P.M., Westphal, H., and Gonzalez, F.J. (1995). Targeted disruption of the alpha isoform of the peroxisome proliferator-activated receptor gene in mice results in abolishment of the pleiotropic effects of peroxisome proliferators. Mol Cell Biol 15, 3012–3022. 10.1128/mcb.15.6.3012

Luukkonen, P.K., Zhou, Y., Sädevirta, S., Leivonen, M., Arola, J., Orešič, M., Hyötyläinen, T., and Yki-Järvinen, H. (2016). Hepatic ceramides dissociate steatosis and insulin resistance in patients with non-alcoholic fatty liver disease. J Hepatol 64, 1167–1175. 10.1016/j.jhep.2016.01.002

Mina, A.I., LeClair, R.A., LeClair, K.B., Cohen, D.E., Lantier, L., and Banks, A.S. (2018). CalR: A web-based analysis tool for indirect calorimetry experiments. Cell Metab 28, 656–666.e651. 10.1016/j.cmet.2018.06.019

Montagner, A., Polizzi, A., Fouché, E., Ducheix, S., Lippi, Y., Lasserre, F., Barquissau, V., Régnier, M., Lukowicz, C., Benhamed, F., et al. (2016). Liver PPARα is crucial for whole-body fatty acid homeostasis and is protective against NAFLD. Gut 65, 1202–1214. 10.1136/gutjnl-2015-310798

Moore, M.P., Cunningham, R.P., Meers, G.M., Johnson, S.A., Wheeler, A.A., Ganga, R.R., Spencer, N.M., Pitt, J.B., Diaz-Arias, A., Swi, A.I.A., et al. (2022). Compromised hepatic mitochondrial fatty acid oxidation and reduced markers of mitochondrial turnover in human NAFLD. Hepatology. 10.1002/hep.32324

Muthusamy, T., Cordes, T., Handzlik, M.K., You, L., Lim, E.W., Gengatharan, J., Pinto, A.F.M., Badur, M.G., Kolar, M.J., Wallace, M., et al. (2020). Serine restriction alters sphingolipid diversity to constrain tumour growth. Nature 586, 790–795. 10.1038/s41586-020-2609-x

Niopek, K., Üstünel, B.E., Seitz, S., Sakurai, M., Zota, A., Mattijssen, F., Wang, X., Sijmonsma, T., Feuchter, Y., Gail, A.M., et al. (2017). A hepatic GAbp-AMPK axis links inflammatory signaling to systemic vascular damage. Cell Rep 20, 1422–1434. 10.1016/j.celrep.2017.07.023

Ochsner, S.A., Abraham, D., Martin, K., Ding, W., McOwiti, A., Kankanamge, W., Wang, Z., Andreano, K., Hamilton, R.A., Chen, Y., et al. (2019). The signaling pathways project, an integrated ‘omics knowledgebase for mammalian cellular signaling pathways. Sci Data 6, 252. 10.1038/s41597-019-0193-4

Oshida, K., Vasani, N., Thomas, R.S., Applegate, D., Rosen, M., Abbott, B., Lau, C., Guo, G., Aleksunes, L.M., Klaassen, C., et al. (2015). Identification of modulators of the nuclear receptor peroxisome proliferator-activated receptor α (PPARα) in a mouse liver gene expression compendium. PLoS One 10, e0112655. 10.1371/journal.pone.0112655

Pang, Z., Chong, J., Zhou, G., de Lima Morais, D.A., Chang, L., Barrette, M., Gauthier, C., Jacques, P., Li, S., and Xia, J. (2021). MetaboAnalyst 5.0: narrowing the gap between raw spectra and functional insights. Nucleic Acids Res 49, W388–w396. 10.1093/nar/gkab382

Park, M., Kaddai, V., Ching, J., Fridianto, K.T., Sieli, R.J., Sugii, S., and Summers, S.A. (2016). A role for ceramides, but not sphingomyelins, as antagonists of insulin signaling and mitochondrial metabolism in C2C12 myotubes. J Biol Chem 291, 23978–23988. 10.1074/jbc.M116.737684

Patel, V.R., Eckel-Mahan, K., Sassone-Corsi, P., and Baldi, P. (2012). CircadiOmics: integrating circadian genomics, transcriptomics, proteomics and metabolomics. Nat Methods 9, 772–773. 10.1038/nmeth.2111

Powers, H.J. (2003). Riboflavin (vitamin B-2) and health. Am J Clin Nutr 77, 1352–1360. 10.1093/ajcn/77.6.1352

Puri, P., Mirshahi, F., Cheung, O., Natarajan, R., Maher, J.W., Kellum, J.M., and Sanyal, A.J. (2008). Activation and dysregulation of the unfolded protein response in nonalcoholic fatty liver disease. Gastroenterology 134, 568–576. 10.1053/j.gastro.2007.10.039

Rinaldo, P., Matern, D., and Bennett, M.J. (2002). Fatty acid oxidation disorders. Annu Rev Physiol 64, 477–502. 10.1146/annurev.physiol.64.082201.154705

Robinson, B.H. (2006). Lactic acidemia and mitochondrial disease. Mol Genet Metab 89, 3–13. 10.1016/j.ymgme.2006.05.015

Rolfe, D.F., and Brown, G.C. (1997). Cellular energy utilization and molecular origin of standard metabolic rate in mammals. Physiol Rev 77, 731–758. 10.1152/physrev.1997.77.3.731

Satapati, S., Kucejova, B., Duarte, J.A., Fletcher, J.A., Reynolds, L., Sunny, N.E., He, T., Nair, L.A., Livingston, K.A., Fu, X., et al. (2015). Mitochondrial metabolism mediates oxidative stress and inflammation in fatty liver. J Clin Invest 125, 4447–4462. 10.1172/jci82204

Scholtes, C., and Giguère, V. (2022). Transcriptional control of energy metabolism by nuclear receptors. Nat Rev Mol Cell Biol. 10.1038/s41580-022-00486-7

Sciacovelli, M., Gonçalves, E., Johnson, T.I., Zecchini, V.R., da Costa, A.S., Gaude, E., Drubbel, A.V., Theobald, S.J., Abbo, S.R., Tran, M.G., et al. (2016). Fumarate is an epigenetic modifier that elicits epithelial-to-mesenchymal transition. Nature 537, 544–547. 10.1038/nature19353

Seo, J., Fortuno, E.S., 3rd, Suh, J.M., Stenesen, D., Tang, W., Parks, E.J., Adams, C.M., Townes, T., and Graff, J.M. (2009). Atf4 regulates obesity, glucose homeostasis, and energy expenditure. Diabetes 58, 2565–2573. 10.2337/db09-0335

Shimano, H., Horton, J.D., Shimomura, I., Hammer, R.E., Brown, M.S., and Goldstein, J.L. (1997). Isoform 1c of sterol regulatory element binding protein is less active than isoform 1a in livers of transgenic mice and in cultured cells. J Clin Invest 99, 846–854. 10.1172/jci119248

Sinal, C.J., Tohkin, M., Miyata, M., Ward, J.M., Lambert, G., and Gonzalez, F.J. (2000). Targeted disruption of the nuclear receptor FXR/BAR impairs bile acid and lipid homeostasis. Cell 102, 731–744. 10.1016/s0092-8674(00)00062-3

Steele, H., Gomez-Duran, A., Pyle, A., Hopton, S., Newman, J., Stefanetti, R.J., Charman, S.J., Parikh, J.D., He, L., Viscomi, C., et al. (2020). Metabolic effects of bezafibrate in mitochondrial disease. EMBO Mol Med 12, e11589. 10.15252/emmm.201911589

Treberg, J.R., Quinlan, C.L., and Brand, M.D. (2011). Evidence for two sites of superoxide production by mitochondrial NADH-ubiquinone oxidoreductase (complex I). J Biol Chem 286, 27103–27110. 10.1074/jbc.M111.252502

Vockley, J., and Ensenauer, R. (2006). Isovaleric acidemia: new aspects of genetic and phenotypic heterogeneity. Am J Med Genet C Semin Med Genet 142c, 95–103. 10.1002/ajmg.c.30089

Ward, P.S., Patel, J., Wise, D.R., Abdel-Wahab, O., Bennett, B.D., Coller, H.A., Cross, J.R., Fantin, V.R., Hedvat, C.V., Perl, A.E., et al. (2010). The common feature of leukemia-associated IDH1 and IDH2 mutations is a neomorphic enzyme activity converting alpha-ketoglutarate to 2-hydroxyglutarate. Cancer Cell 17, 225–234. 10.1016/j.ccr.2010.01.020

Waskowicz, L.R., Zhou, J., Landau, D.J., Brooks, E.D., Lim, A., Yavarow, Z.A., Kudo, T., Zhang, H., Wu, Y., Grant, S., et al. (2019). Bezafibrate induces autophagy and improves hepatic lipid metabolism in glycogen storage disease type Ia. Hum Mol Genet 28, 143–154. 10.1093/hmg/ddy343

Xu, Y., Zhu, Y., Hu, S., Xu, Y., Stroup, D., Pan, X., Bawa, F.C., Chen, S., Gopoju, R., Yin, L., et al. (2021). Hepatocyte nuclear factor 4α prevents the steatosis-to-NASH progression by regulating p53 and bile acid signaling (in mice). Hepatology 73, 2251–2265. 10.1002/hep.31604

Yang, L., Garcia Canaveras, J.C., Chen, Z., Wang, L., Liang, L., Jang, C., Mayr, J.A., Zhang, Z., Ghergurovich, J.M., Zhan, L., et al. (2020). Serine catabolism feeds NADH when respiration is impaired. Cell Metab 31, 809–821.e806. 10.1016/j.cmet.2020.02.017

Yavarow, Z.A., Kang, H.R., Waskowicz, L.R., Bay, B.H., Young, S.P., Yen, P.M., and Koeberl, D.D. (2020). Fenofibrate rapidly decreases hepatic lipid and glycogen storage in neonatal mice with glycogen storage disease type Ia. Hum Mol Genet 29, 286–294. 10.1093/hmg/ddz290

Ye, J., Mancuso, A., Tong, X., Ward, P.S., Fan, J., Rabinowitz, J.D., and Thompson, C.B. (2012). Pyruvate kinase M2 promotes de novo serine synthesis to sustain mTORC1 activity and cell proliferation. Proc Natl Acad Sci U S A 109, 6904–6909. 10.1073/pnas.1204176109

